# Active learning for improving out-of-distribution lab-in-the-loop experimental design

**DOI:** 10.1101/2025.02.26.640110

**Authors:** Daria Balashova, Robert Frank, Svetlana Kuzyakina, Dominique Weltevreden, Philippe A. Robert, Geir Kjetil Sandve, Victor Greiff

## Abstract

The accurate prediction of antibody-antigen binding is crucial for developing antibody-based therapeutics and advancing immunological research. Library-on-library approaches, where many antigens are probed against many antibodies, can identify specific interacting pairs. Machine learning models can predict target binding by analyzing many-to-many relationships between antibodies and antigens. However, these models face challenges when predicting interactions when test antibodies and antigens are not represented in the training data, a scenario known as out-of-distribution prediction. Generating experimental binding data is costly, limiting the availability of comprehensive datasets. Active learning can reduce costs by starting with a small labeled subset of data and iteratively expanding the labeled dataset. Few active learning approaches are available to handle data with many-to-many relationships as, for example, obtained from library-on-library screening approaches. In this study, we developed and evaluated fourteen novel active learning strategies for antibody-antigen binding prediction in a library-on-library setting and evaluated their out-of-distribution performance using the Absolut! simulation framework. We found that three of the fourteen algorithms tested significantly outperformed the baseline where random data are iteratively labeled. The best algorithm reduced the number of required antigen mutant variants by up to 35%, and sped up the learning process by 28 steps compared to the random baseline. These findings demonstrate that active learning can improve experimental efficiency in a library-on-library setting and advance antibody-antigen binding prediction.

## Introduction

Adaptive immunity is a specialized branch of the immune system that provides long-lasting and highly specific defense against pathogens. It mainly consists of T cells and B cells, which work together to recognize and respond to specific antigens (Ags) thanks to their highly diverse set of T- and B-cell receptors (BCRs and TCRs), respectively (Murphy and Weaver, 2017). In this manuscript, we focus on antibodies (Abs) – proteins produced by B cells that bind to antigens with high specificity to neutralize pathogens or mark them for further elimination. The ability of B cells to generate a diverse array of Abs is facilitated through processes such as gene rearrangement, p/n-nucleotide addition, and somatic hypermutation, enabling them to effectively target a wide range of pathogens (Murphy and Weaver, 2017; Tonegawa, 1983).

Monoclonal Abs (mAbs) have become increasingly significant in recent years, particularly in the fields of oncology and immunology. Their market value is projected to reach 445 billion USD by 2028 (Lyu et al., 2022). Due to their ability to bind specific targets while being well tolerated, mAbs are not only essential in therapeutic applications but also valuable as probes in laboratory research (Nelson, 2000).

Recently, their relevance was underscored during the SARS-CoV-2 pandemic, where therapeutics mAbs could confer protection against infection (Pochtovyi et al., 2023). However, the rapid mutation rate of the SARS-CoV-2 virus often rendered available Abs ineffective, particularly due to changes in the receptor-binding domain of the Spike protein on the virus surface (Ragonnet-Cronin et al., 2023). Even when an Ab becomes ineffective against a viral variant, other Ab cocktails might bind effectively to the new Ag variant (Taft et al., 2022; Ehling et al., 2024).

The prediction of Ab-Ag binding for novel sequences is pivotal in the development of new monoclonal Ab-based therapeutics and in enhancing fundamental research (Figure 1A) (Graves et al., 2020; O’Donnell et al., 2024). Several machine learning (ML) based tools have recently been developed to predict Ab-Ag binding across Abs and Ags. AbAgIntPre, a deep learning method, predicts Ab-Ag interactions based solely on amino acid sequences, achieving an ROC-AUC of 0.82 (Huang et al., 2022). AttABseq, an attention-based model, excels in predicting binding affinity changes due to mutations, outperforming other sequence-based models by 120% (Jin et al., 2024). Additionally, AntBO, a Bayesian optimization framework, efficiently designs CDRH3 sequences with high affinity, outperforming genetic algorithms and reducing experimental iterations (Khan et al., 2023). These advancements are transforming antibody design by enabling faster, cost-effective predictions. Chinery et al. employed computational methodologies, including BLOSUM (Henikoff and Henikoff, 1992), AbLang (Olsen et al., 2022), ESM (Lin et al., 2023), and Protein-MPNN (Dauparas et al., 2022), to design diverse antibody libraries derived from a Trastuzumab sequence (Chinery et al., 2024).

**Figure 1:**
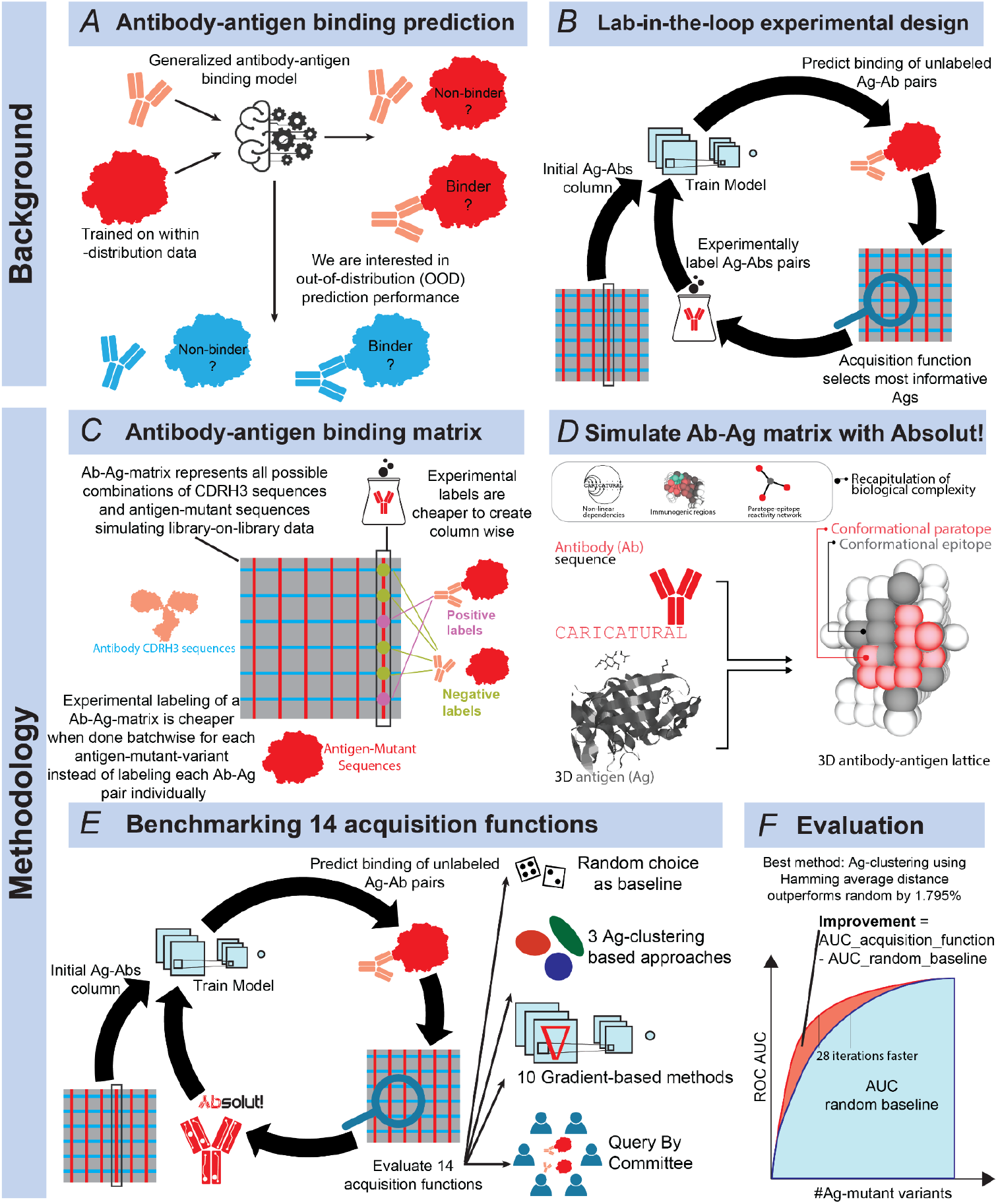
Enhancing data-efficient training of generalized antibody-antigen binding prediction models through Active Learning approaches. (A) The precise out-of-distribution (OOD) prediction of Ab-Ag binding, particularly for novel Ags, is crucial for the advancement of monoclonal Ab-based therapeutics and foundational research. While Ab-Ag binding classification can be theoretically conducted for all possible Ab-Ag pair combinations, it is impractical to experimentally label every pair due to resource constraints; hence, only a selective subset can be annotated. Generalized OOD Ab-Ag binding prediction aids in inferring the binding potential of unlabeled Ab-Ag pairs. (B) Active learning (AL) techniques are utilized to select new Ag-related batches for labeling, thus maximizing their informative value based on existing labeled data. This process creates a dynamic interaction between the AL algorithm and experimental wet-lab Ab batch labeling (lab-in-the-loop). Starting with an initial set of Ags, this iterative cycle includes model training, querying the most informative unlabeled Ags (batches of Ag-Abs pairs where the Ag is the same for the entire batch), experimental labeling, and dataset enrichment, with the refined dataset used for successive training. Efficient labeling of Ab-Ag pairs involves the inclusion of an Ag and multiple randomly mutated CDRH3 sequences. (C) Library-on-library data is organized into a two-dimensional Ab-Ag-matrix, where each potential Ab and Ag-mutant-variant sequence pair is associated with a label. Experimentally labeling the matrix is expensive. It is cheaper to label entire batches (libraries) of Abs against a single Ag mutant variant than to label Ab-Ag-mutant-variant pairs independently. We precompute an extensive data matrix to function as an oracle in lab-in-the-loop simulations, enabling us to emulate smaller, labeled Ab-Ag matrices generated by an AL algorithm. (D) The Absolut! framework calculates the binding affinity between CDRH3 sequences and Ags by discretizing the Ag’s structural conformation and exhaustively evaluating all potential binding configurations for the CDR sequence. We introduce mutations at a binding hotspot on an Ag and compute the binding affinities of a set of Abs against these Ag mutant variants to create a simulated Ab-Ag binding matrix. (E) We simulate lab-in-the-loop training on Ab-Ag matrices generated using the Absolut! framework. Fourteen different acquisition functions are assessed against a uniform random acquisition function baseline, each tested with 200 distinct random initializations. (F) The efficacy of each AL strategy is examined by comparing the area under the Ag-iterations/ROC AUC curve for each acquisition function against the baseline curve area provided by the random acquisition function. The most effective acquisition function is the Ag sequence clustering strategy using the average Hamming distance, which surpasses the random strategy by 1.795%.

To develop ML methods that perform effectively across a wide range of Ab-Ag interactions, it is essential to have a diverse array of labeled data. The ability to generalize successfully to new sequences heavily relies on the specific sequences for which labeled data are available. Given the high cost associated with labeling Ab-Ag sequences, identifying the optimal set of sequence combinations for labeling is of paramount importance. Due to the high costs associated with experimentally labeling Ab-Ag binding sequence pairs (Engelhart et al., 2022a), it is essential to meticulously design experimental plans to optimize resource utilization. An effective strategy in this context is the lab-in-the-loop approach (Buntz, 2024). This method involves the iterative labeling of batches of Abs against a single Ag, followed by model training on the collected data. Subsequently, an acquisition function is employed to identify the next most informative batch of Ab-Ag pairs for labeling (Figure 1B).

Active learning (AL) is a machine learning approach where the model selectively queries an information source (such as a human expert or an oracle) to label new data points, rather than passively learning from a fixed dataset (Cohn et al., 1994; Ren et al., 2021; Settles, 2009). Therefore, it optimizes data selection for labeling, which is particularly useful in scenarios where labeled data is scarce or expensive to obtain. In the context of Ab-Ag binding, AL can be used to minimize the number of lab tests required by acquiring smaller datasets iteratively instead of a single large initial random dataset, while maximizing model performance. The learner, trained on a small labeled dataset, selects the most informative instances from an unlabeled pool to be tested in the lab. Experimental results then serve as feedback, refining the model and improving its predictive performance while reducing the overall experimental workload.

AL techniques have been employed to predict Ab sequences binding to their target Ags with high efficacy despite using small datasets, successfully identifying at least nine novel heavy chain complementarity-determinant region Ab sequences per Ag from a dataset of only six-thousand examples (Seo et al., 2022). Additionally, another AL framework has been developed to iteratively propose promising Ab candidates for evaluation, significantly accelerating the identification of improved binders and optimizing the surrogate modeling approach, as demonstrated through both pre-computed data pools and realistic full-loop scenarios (Gessner et al., 2024).

Technically speaking, several strategies have been proposed for ML models to propose the next datasets to generate. A pivotal strategy in AL involves selecting data points that maximize a model’s uncertainty. In the Query-by-Committee (QBC) approach, multiple models are trained as committee members, and the data instance that generates the greatest disagreement among them is chosen for labeling (Seung et al., 1992). A different approach to measuring uncertainty leverages model gradients – large gradients indicate high uncertainty, making such instances valuable for refinement (Lee and AlRegib, 2020). The BADGE algorithm (Ash et al., 2019) combines gradient-based uncertainty with sample diversity, improving sample selection. Alternatively, diversity-based approaches emphasize selecting data that spans a wide range of the input space, ensuring the model learns from a varied set of examples (Sener and Savarese, 2018). This strategy focuses on the inherent characteristics of the data rather than the model’s internal state. While model-based approaches, such as QBC or gradient-based selection, rely on the model’s internal state to guide data acquisition, diversity-based approaches prioritize the distribution of data itself.

The training of ML models that predict the binding of Ab-Ag sequences ideally requires experimental datasets that label a set of many Ab sequences to many Ag sequences. The generation of experimental data entails the development of an Abs library, which is subsequently screened against libraries of Ags. AlphaSeq (Engelhart et al., 2022b) is a library-on-library technology that enables the simultaneous measurement of millions of protein-protein interactions. This is achieved by genetically reprogramming yeast cells to combine their genetic material upon interaction, with next-generation sequencing subsequently quantifying these events. This approach allows for the assessment of specificity and multi-target affinity across extensive libraries of Abs and Ags in a single experiment. Consequently, the Ab-Ag binding data can be represented as a two-dimensional matrix, where each Ag can be screened for interaction with all possible Abs and vice versa (Figure 1 C). Interactions can be binary (binding or non-binding), semi-quantitative (non-binder, low, medium, high affinity), or quantitative (affinity measurements).

The scarcity of large lab-tested datasets often necessitates the use of synthetic or publicly available data that are heterogeneous in terms of experimental technique and affinity thresholds to define binding pairs. To tackle the complexities of Ab-Ag interactions and simulate realistic experimental conditions at a large-scale, the Absolut! simulation framework was developed (Robert et al., 2022). Absolut! enables the generation of synthetic library-on-library binding datasets by simulating the binding process of Ab-Ag binding at a molecular level in a deterministic fashion and with the same settings across many Ab and Ags. This framework creates a controlled environment where different Ags and Abs can be systematically tested against each other, producing data that mimics real-world experimental library-on-library outcomes (Figure 1 D). Absolut! has been used to develop optimization techniques for Ab design (Khan et al., 2023; Maraval et al., 2023; Towers et al., 2024). Here, we generated a dataset consisting of 117 Ags and 2230 Abs resulting in 260 910 binding pairs, using Absolut!, providing a robust foundation for training and evaluating machine learning models. We precompute the data to serve as an oracle in lab-in-the-loop simulations. By simulating various Ag variants and their interactions with Abs, Absolut! helps to explore the vast sequence space efficiently and supports the development of more accurate predictive models.

In this study, we explore the potential of AL techniques to enhance the selection and sequencing of Ags in iterative laboratory experiments, aiming to reduce the number of experiments needed to accurately predict Ab-Ag binding, which is crucial for efficient therapeutic antibody development and immune research. We utilize simulations of lab-in-the-loop experiments with Absolut! to perform this exploration of AL techniques (Figure 1E), specifically focusing on library-on-library datasets, where many-to-many Ab-Ag interactions are systematically tested. The objective is to reach the necessary accuracy in Ab-Ag binding prediction models with fewer experimental iterations, compared to the random addition of data, across various testing conditions specific to Abs and Ags. Also, the capacity to compare the performance of different ML strategies on large, fast and cheap simulated datasets is critical to know in advance which strategy will work best on experimental datasets. We hypothesize that these techniques will allow the model to achieve the desired performance level more efficiently, thereby reducing the time and cost associated with experimental Ab-Ag binding studies. Performance enhancement is assessed by integrating the ROC AUC across the number of iterations and comparing it against a baseline (Figure 1F).

## Results

We compared a range of active learning (AL) strategies to identify the most informative antibody-antigen (Ab-Ag) pairs for labeling in a resource-efficient manner. These strategies included both model-based and diversity-based approaches, each focusing on maximizing the informativeness of selected pairs. We differentiate between AL strategies that use trained models to estimate the usefulness of unobserved Ag mutant variants (Model-based), and strategies that only operate on the sequences (Diversity-based).

### Model-based strategies

A pivotal strategy in AL involves selecting data points that maximize a model’s uncertainty. We implemented two primary model-based strategies: Query-by-Committee (QBC) and Gradient-Based Uncertainty.

In the QBC approach, multiple models are trained as committee members, and the data instance that generates the greatest disagreement among them is chosen for labeling (Seung et al., 1992). We trained a committee of five convolutional neural networks, where higher variance across model predictions indicated greater disagreement. We identified new Ags for labeling by focusing on those that generated the highest disagreement among committee members.

An alternative measure of uncertainty leverages the model’s gradient, given that optimization of neural network parameters typically involves gradient descent (Ash et al., 2019; Lee and AlRegib, 2020). If a model exhibits a large gradient for a particular instance, it means that the loss function is highly sensitive to changes in the weights for that instance. This results in a substantial update to the weights during gradient descent. Therefore, the gradient norm as an indicator of uncertainty allows the separation of instances that can be predicted confidently from those that remain uncertain, and thus prioritizing the uncertain Ags for additional experimental measurements.

### Diversity-based strategy

Diversity-based approaches emphasize selecting data that spans a wide range of the input space, ensuring the model learns from a varied set of examples (Sener and Savarese, 2018). This strategy focuses on the inherent characteristics of the data rather than the model’s internal state. Diversity-based approaches rely on the intuitive assumption that subjects located near each other in a given metric space share common properties, making them equally informative for the model while also introducing redundancy into the dataset. The next Ag was chosen based on its distance from the training set, considering both the similarity metric and the method for computing distance. We explored Hamming distance and alignment-based distance as similarity metrics. For Hamming distance, we applied two selection strategies: average distance, where the Ag with the largest mean Hamming distance to all labeled Ags was selected to maximize overall diversity, and minimum distance, where the Ag with the largest smallest Hamming distance to any labeled Ag was chosen to ensure it was not overly similar to its closest counterpart. For alignment-based distance, only the average distance strategy was used.

### Simulation-based evaluation of performance

As a baseline for comparison, we implemented a random selection strategy where binding data between a randomly selected Ag and all Abs was iteratively added to the training set. This approach provided a neutral reference point, allowing us to assess the effectiveness of our AL strategies in selecting the most informative Ags. By comparing AL methods against this baseline, we could determine whether targeted selection led to improved predictive performance over purely random sampling.

The Ab-Ag matrix represents interactions between antigen sequences and 11-mer sliding windows of antibody CDRH3 sequences, where each Ab sequence is broken down into overlapping 11-mer segments. In our evaluation, we assessed each AL strategy against a baseline random selection approach using synthetic Ab-Ag matrices generated within the Absolut! simulation framework. The task was framed as a binary classification problem, where the model predicted the binding or non-binding status of Ab-Ag pairs. Performance was measured by applying the AL approach and evaluating the model at each iteration. At every iteration, the model was trained on the Ab-Ag binding data available up to that point and then used to predict the binding or non-binding status of Ab-Ag pairs. Based on these predictions and the true labels, we calculated the receiver operating characteristic area under the curve (ROC AUC) on the test dataset. For each iteration (x-axis), the model’s performance was recorded as ROC AUC (y-axis), forming an active learning curve (ALC). The area under ALC was then used as the final performance metric for the AL strategy. Finally, we compared the performance of each AL approach to that of the random baseline to assess its effectiveness.

The training dataset grew with each iteration, adding binding data for one Ag mutant variant (with 1-3 mutations from initial Ag) selected randomly or by an active learning strategy with all Abs at every step. The test datasets remained fixed, ensuring consistency across iterations. The experiment included three distinct test datasets, each designed to evaluate the models’ performance under different conditions of sequence novelty. These datasets were designed based on the two-dimensional structure of the data, where Ag sequences were split into 80% training and 20% testing, while Ab sequences were split evenly into 50% training and 50% testing. The simultaneous split across both dimensions (Ab-Ag) guarantees no data leakage.

The first dataset, referred to as the Test set (Figure 3 A), featured entirely unseen Ab and Ag sequences during testing, creating a stringent out-of-distribution (OOD) scenario, and included 11 339 Ab-Ag pairs. The second dataset, TestSharedAG (Figure 3 B), contained 46 342 Ab-Ag pairs with novel Ab sequences but included Ag sequences that were familiar to the model, allowing for a partial reliance on known Ags during prediction. The final dataset, TestSharedAB (Figure 3 C), was structured with 11 316 Ab-Ag pairs consisting only novel Ag sequences, while the Ab sequences were previously encountered during training. The training dataset contained 46 248 Ab-Ag pairs (492 Abs and 94 Ags). This setup provided a graded assessment of each AL strategy’s robustness and effectiveness under different OOD conditions.

The Hamming Average Distance approach achieves the highest performance gain in the Test set (Figure 2 A), with a significant improvement (Bonferroni correction applied) of 1.795% over the baseline. It is particularly effective in settings with 20-70 Ags. While the random selection method requires 54 Ags, AL achieves the same accuracy with only 35 Ags, reducing the required number by 35%. The final accuracy of the random selection model is achieved in 66 steps using the Hamming Average Distance, reducing the number of steps by 28. QBC also enhances performance in the Test set, providing a 0.766% gain over the baseline. This highlights the suitability of diversity based approaches for optimizing performance in Ab-Ag binding prediction tasks in a complete out-of-distribution scenario where neither Ab nor Ags were known in the prediction task.

**Figure 2:**
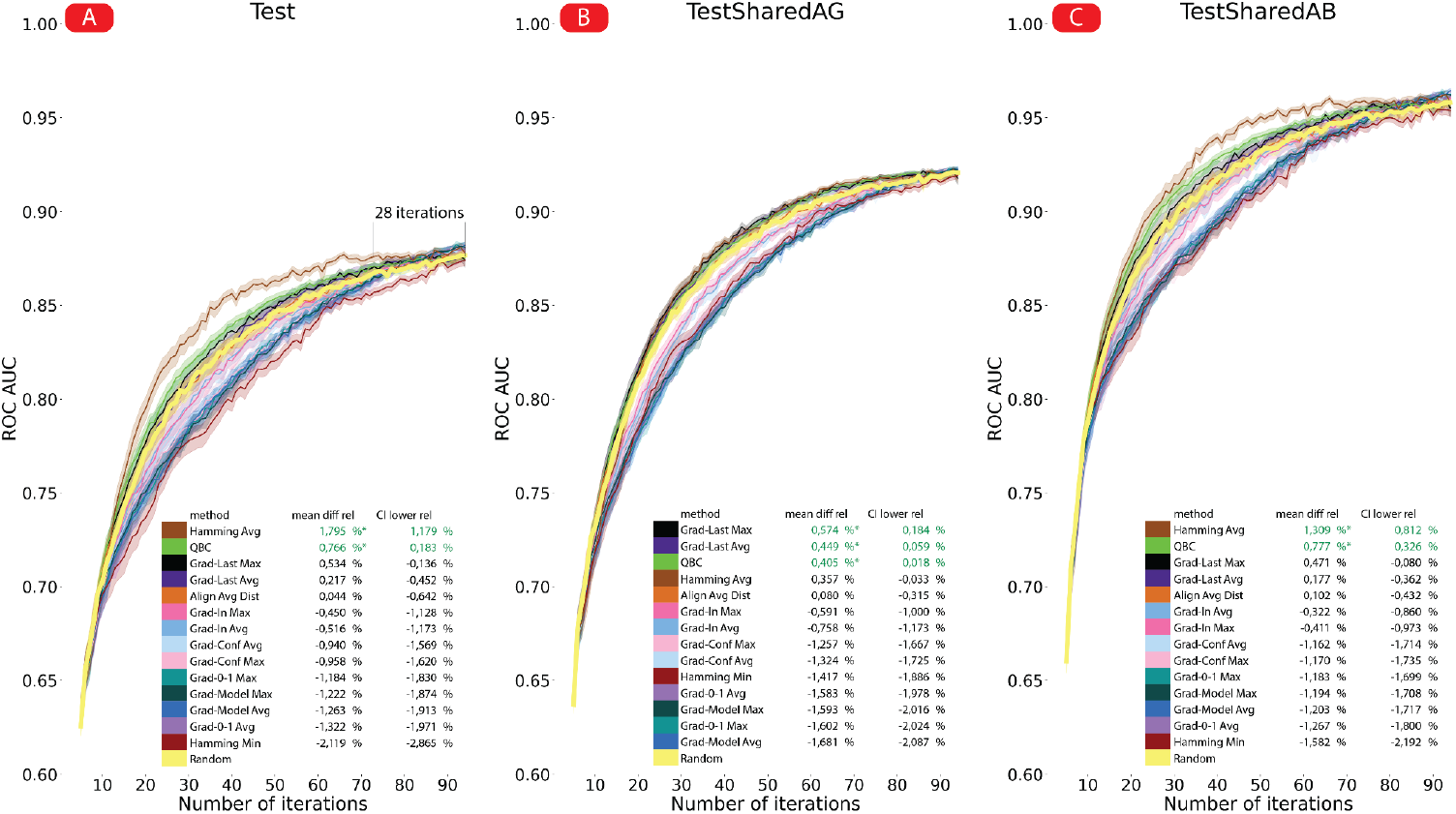
Three out of fourteen active learning acquisition functions significantly outperform the random baseline. The performance of fourteen different active learning (AL) acquisition functions is shown on three different dataset (A) Test: Ab-Ag binding prediction where neither the Ab nor the Ag was available during training. (B) TestSharedAG: Ab-Ag binding prediction where the Ags were available during training. (C) TestSharedAB: Ab-Ag binding prediction where the Abs were available during training). Each data point on the graph represents the performance (ROC AUC) of a single model trained on a sequence of Ags (the number of Ags is indicated on the x-axis) chosen by a particular AL method. Each AL method, along with the random baseline, was evaluated over 200 different random repetitions of the experiment. Box plots illustrating the distribution of these experiments can be found in Supplementary Figures 1, 2, and 3. The best methods are: Hamming Average Distance (1.795% improvement on Test, reducing the number of required Ag mutant variants by up to 28, and 1.309% improvement on TestSharedAB) and “Gradient on Last Layer Max” (0.574% improvement on TestSharedAG). Full statistics can be found in supplementary table 1.

In the TestSharedAG set (Figure 2 B) ROC AUC values ranging from 0.64 to 0.92. QBC leads to a 0.405% improvement over the random baseline, while Gradient-Based uncertainty methods, utilizing the gradient norm in the model’s last layer, yield increases of 0.574% and 0.449% for Maximum and Average norms, respectively.

For the TestSharedAB set (Figure 2 C), the Hamming Average Distance approach again demonstrates strong performance, achieving a 1.309% improvement over the baseline. QBC provides a 0.777% gain, while Gradient-Based uncertainty methods show smaller yet consistent improvements. The overall performance in TestSharedAB is intermediate between Test and TestSharedAG, with ROC AUC values ranging from approximately 0.66 to 0.96. These findings indicate that AL methods effectively enhance prediction accuracy for this dataset, though the degree of improvement varies by method and dataset characteristics.

## Discussion

This study introduced and evaluated fourteen active learning (AL) strategies designed for predicting antibody-antigen (Ab-Ag) binding, using the Absolut! simulation framework (Robert et al., 2022) to emulate a data acquisition context suitable for evaluating AL strategies. Three strategies – Hamming Average Distance, Gradient-Based uncertainty (Last Layer Max) and Query-by-Committee – demonstrated significant performance gains over the random baseline. The Hamming Average Distance method achieved the highest improvement, with a 1.795% increase in the area under the active learning curve compared to the random selection baseline in the Test dataset, reducing the required number of Ag mutant variants by 35% and achieving accuracy comparable to the last step of random selection model 28 steps earlier. Overall, the AL strategies made data use much more efficient, significantly cutting down the need for experimental labeling.

The results show that tailored AL methods are very effective at improving how Ab-Ag pairs are chosen for labeling. The Hamming Average Distance method performed especially well, showing that selecting diverse Ags based on sequence differences (i.e., a diversity-based approach) leads to better Ag selection and improved model performance. Notably, this method can be implemented without relying on the AL paradigm. Instead, clustering-based approaches like the average Hamming distance can be applied at the outset, enabling the selection of Ag mutant variants before iterative learning begins. Collectively, the proposed strategies represent powerful options for addressing the inherent challenges of library-on-library Ab-Ag binding prediction, particularly in out-of-distribution scenarios.

Several limitations warrant consideration. The study focused on binary classification of Ab-Ag binding, which may not capture the nuanced differences in binding affinity required for therapeutic applications. The Ag mutations studied were limited to 1-3 alterations from a base sequence, which may not generalize to highly diverse or unrelated Ags. Additionally, while Absolut! provides a robust simulation framework, it may not fully replicate the complexities of real-world Ab-Ag interactions, including structural and biochemical factors that influence binding. Finally, the experimental datasets were constrained to specific scenarios, and broader validation is necessary to assess the performance of these strategies across a wider range of Abs and Ags. The evaluation across three different test datasets provided key insights into how AL strategies generalize under varying levels of sequence novelty on the Ab or Ag side. Further analysis of Ab and Ag similarity between training and test sets could refine AL strategies by uncovering patterns in generalization.

Future research could also explore regression-based AL strategies to predict binding affinities on a continuous scale, providing more detailed insights into Ab-Ag interactions (Li et al., 2023). Further improvement of AL performance could be achieved by applying diversity and model based approaches simultaneously. Expanding the scope of the dataset to include a greater diversity of Abs and Ags, such as unrelated Abs or Ag mutant variants, would improve model versatility (Jin et al., 2024) and might lead to an even greater AL performance than diversity based or model based AL methods on their own. Moreover, integrating interpretability techniques, such as SHAP (Lundberg and Lee, 2017) or LIME (Ribeiro et al., 2016), could enhance understanding of model predictions and identify potential biases in training data.

The model-based approaches assessed in this study are predicated on the assumption that model uncertainty correlates with high informational value of the unlabeled Ab-Ag sequence pairs. Although our findings indicate that this assumption yields beneficial results, calculating the expected improvement directly could potentially enhance the efficiency of AL strategies. This method, known as Bayesian experimental design, is typically computationally intensive and necessitates approximations, such as those achieved via policy networks (Rainforth et al., 2023) or Monte Carlo approximations.

We advocate for the use of AL for antigen mutation selection rather than random sampling when creating Ab-Ag matrices. However, experimental validation of the proposed AL strategies using real-world Ab-Ag datasets is crucial to confirm their practical utility. To determine which AL method performs best under different out-of-distribution conditions, additional testing on experimental data is needed. Large real-world Ab-Ag datasets can be used to optimize AL methods and select the most effective ones, ensuring their applicability (Engelhart et al., 2022b; Mason and Reddy, 2024). This, in turn, will facilitate more efficient data acquisition, reducing experimental costs while maintaining predictive accuracy. Achieving this requires close interdisciplinary collaboration between computational and experimental researchers to bridge the gap between predictive modeling and therapeutic application.

## Methods

### 1 Data generation

The antibody-antigen (Ab-Ag) binding data utilized in this study was generated using the 3D simulation framework Absolut! (Robert et al., 2022). Machine learning techniques require substantial amounts of labeled data for training and validation; however, acquiring labeled Ab-Ag data is challenging and costly, necessitating the use of simulated datasets. The datasets created by Absolut! are designed to mimic the noise and principles observed in real-world Ab-Ag binding data. It has been demonstrated that the performance of machine learning methods is comparable on datasets generated by Absolut! and experimental data (Robert et al., 2022): while Absolut! does not directly predict real-world Ab-Ag binding, it facilitates the development of machine learning strategies that, when effective on simulated data, also perform well with experimental data. (Robert et al., 2022) showed that improvements in machine learning predictions for simulated data similarly enhanced predictions for real-world data.

Absolut! constructs a discretized lattice representation of a protein-antigen from the Protein Data Bank (Robert et al., 2022; Burley et al., 2023). Based on this lattice, it computes the optimal binding for the Ab sequence, specifically the CDRH3. The CDRH3 sequences, sourced from mice, were capped at 11 amino acids; for sequences longer than 11, Absolut! estimated the binding energy for 11-mer segments. In this research, all CDRH3 sequences were tested against antigen mutant variants for a single antigen, 1ADQ_A (Corper et al., 1997). The CDRH3 sequences in our dataset were selected by filtering all CDRH3 sequences from the Absolut! dataset through two steps. Firstly, sequences were filtered based on their binding affinity with the 1ADQ_A antigen, selecting the top 1% with the best binding scores. Secondly, sequences were clustered into groups (binding hotspots) that interact with the same four amino acids (or more) on the Ag, with each sequence belonging to only one cluster. The largest CDRH3 sequence cluster (hotspot) was then chosen, and the four defining amino acids on the antigen were termed the binding hotspot.

The four amino acids in the binding hotspot were systematically mutated, with up to 3 amino acids being modified. This involved 80 one-point mutations (covering all possible single-residue changes), 500 randomly sampled two-point mutations, and 1,500 randomly sampled three-point mutations, culminating in 2,080 antigen mutations in total. It is assumed these mutations do not significantly alter the protein structure of the antigen. The final dataset comprised 2,080 antigen mutations of 1ADQ and 2,230 antibodies.

To increase the dataset’s difficulty and highlight the utility of active learning (AL) approaches, specific criteria were applied to refine the raw dataset, initially consisting of randomly sampled antigen mutations. All overlapping three-point mutations were removed, as were two-point mutations contained within three-point mutations. Furthermore, all antibody 11-mers used were derived from different antibodies, ensuring only the best 11-mer segment from each antibody was included, excluding any other segments from the same source. Consequently, the final dataset contained 117 Ag mutant variants and 2,230 Abs.

The data was subsequently divided into training and testing datasets, with Ag sequences allocated in an 80%/20% ratio and Ab sequences in a 50%/50% ratio. Owing to the data’s two-dimensional structure, three distinct test datasets were formulated (Figure 3): TestSharedAG, which utilizes Ag mutant variants from the training dataset and Ab sequences from the test dataset; TestSharedAB, which contains Ag mutant variants from the test dataset and Ab sequences from the training dataset; and Test, which comprises both Ag mutant variants and Ab sequences from the test dataset.

**Figure 3:**
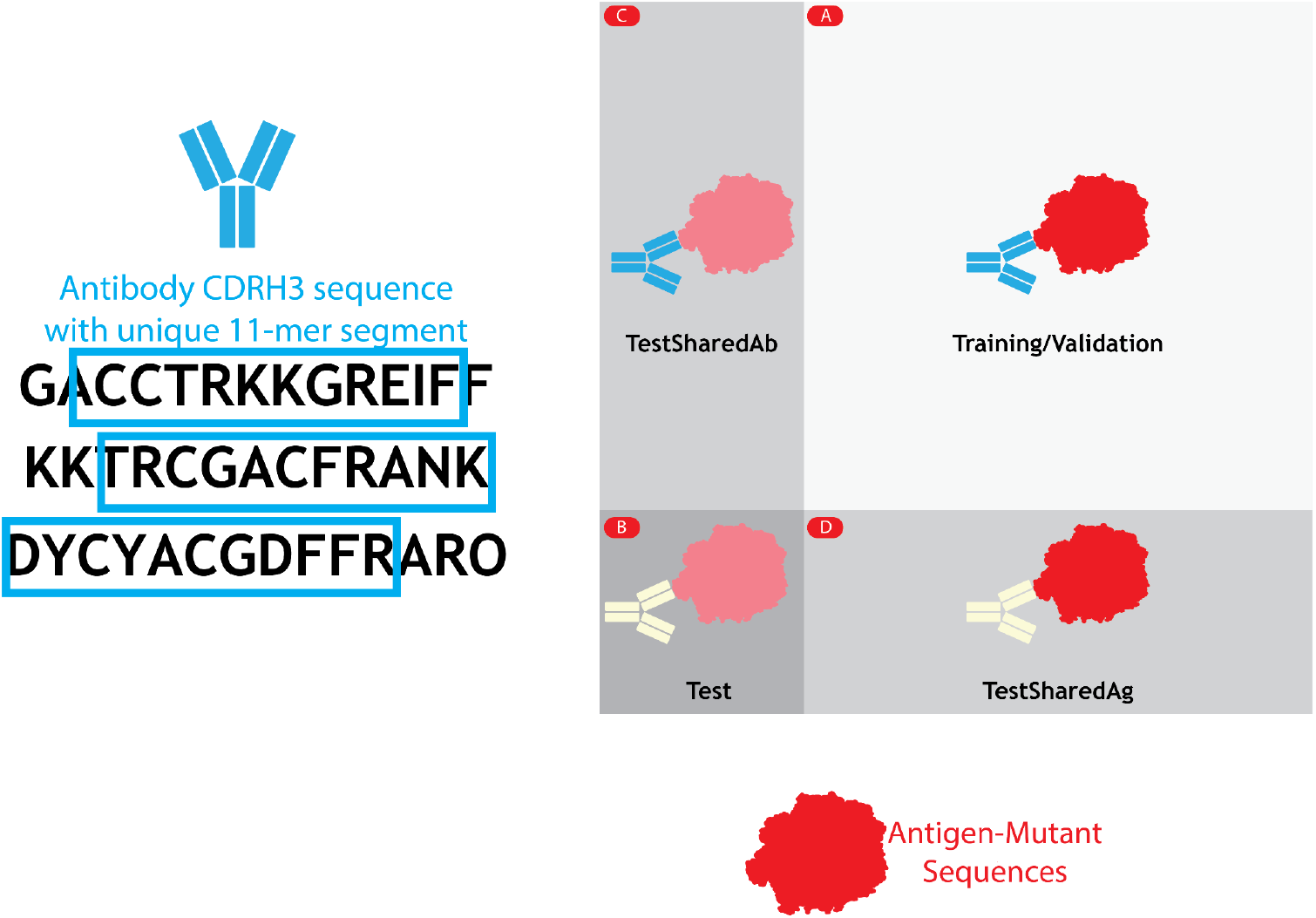
The Ab-Ag-matrix illustrates the 2D structure of our dataset, demonstrating how Abs and Ags are divided into training and test datasets. The x-axis represents the sequences of different Ag mutant variants, while the y-axis corresponds to the CDRH3 sequences. For each possible combination of CDRH3 sequences and Ag mutant variant sequences, we compute the binding affinity for all 11-mer segments of CDRH3 using Absolut!, selecting the one with the highest binding affinity for the CDRH3. (A) The Ab-Ag-matrix is segmented along both dimensions to create a training/validation dataset and a (B) testing dataset, ensuring no overlap of Ag mutant sequences or CDRH3 sequences between them. Additionally, two minor datasets are designed: (C) the TestSharedAB dataset, which comprises Ag mutant variant sequences different from those in the training dataset and CDRH3 sequences identical to those in the training dataset; and (D) the TestSharedAG dataset, which includes CDRH3 sequences distinct from those in the training dataset and Ag mutant variants that are identical to those in the training dataset.

### 2 Active learning approaches

#### 2.1 Model/uncertainty based

The process begins with a small initial training set of Ab-Ag pairs. In each iteration, the model evaluates the uncertainty of potential Ags by measuring the variance or gradients of predictions for Ab-Ag pairs, highlighting the Ags with the greatest model uncertainty. The most uncertain Ags are selected for inclusion in the training set. These Ags are paired with Abs and added to the training data, after which the model is retrained. This cycle repeats, with the model refining its predictions as more Ags are added.

##### 2.1.1 Query-by-Committee

###### Approach

The central idea of **Query-by-Committee** (QBC) AL strategy is to create a committee of models and to select new data points that cause the greatest disagreement among these models (Seung et al., 1992).

###### Neural network

In this study, the committee consists of five convolutional neural networks, each trained independently on a randomly selected 50% subset of the available labeled data. This sampling strategy ensures diversity in the training data for each model, which is essential for generating varying predictions on the same data instances. The committee size is set to five, balancing computational efficiency with the need for robust disagreement measurement.

###### AL acquisition

**Step 1:** Select candidate Ab-Ag pairs where the Ags are not yet part of the training set.

**Step 2:** Predict outcome and measure variance. Each model in the committee generates predictions for these candidate Ab-Ag pairs. The variance of the predictions across the models is then computed for each Ag.

**Step 3:** Quantify disagreement. For each Ag, we predict binding probabilities for all paired Abs using five models and compute the variance of these predictions for each Ab-Ag pair. Since each Ag is paired with multiple Abs, this results in a distribution of variance values for each Ag.

To quantify disagreement, we take the 0.9 quantile of this variance distribution, providing a robust measure of uncertainty for each Ag.

**Step 4:** Select Ag. The Ag with the highest disagreement score is selected for inclusion in the next iteration of the training set.

###### Experimental Setup

The initial model was trained using a base set of five Ags. The AL process is repeated for 100 iterations, with new Ags added to the training set based on the QBC-driven selection.

##### 2.1.2 Gradient-Based uncertainty

Previously, the use of gradient norm as a measure of uncertainty was proposed by several authors, who implemented approaches based on the gradient of the loss (Ash et al., 2019), including confounding labels approach (Lee and AlRegib, 2020). In our research we implemented a variety of gradient-based methods, including classical gradient with respect to the model’s last layer for predicted labels and confounding labels. Our methods can be separated into two groups: methods that use different loss functions (including different put labels) and methods that vary parameters, with respect to which gradient is counted.

The first method – **Gradient on Last Layer Average** (Grad-Last Avg) uses the gradient of the loss function with respect to the last linear layer in the model as a measure of uncertainty. The implementation algorithm of the method includes the following steps:

**Step 1:** Define the initial training set.

**Step 2:** Train the model on the initial training set.

**Step 3:** For each Ag not included in the initial training set:

1. Calculate the model prediction for all Abs and the given Ag;
2. Calculate the value of the loss function based on the model prediction for all Abs and the given Ag;
3. Calculate the gradient of the loss function with respect to the last linear layer of the model and then calculate its norm for the given Ag and all Abs;
4. Calculate the average value of the gradient norm over all Abs for the given Ag.

**Step 4:** Add the Ag with the largest average gradient norm over all Abs to the training set.

The **Gradient on Last Layer Max** (Grad-Last Max) method differs from the Gradient on Last Layer Average method only in using the maximum gradient norm among all Abs instead of the average value of the gradient norm for all Abs (for a fixed Ag).

The second method – **Gradient 0-1 Average** (Grad 0-1 Avg) replaces the true model predictions for each Ab-Ag pair by first substituting 0 into the loss function (which corresponds to an Ag not bound to any Ab) for all Ab-Ag pairs and calculating its gradient, and then substituting 1 (which corresponds to an Ag bound to all Abs). The method tries to estimate how different the changes in the model parameters are for two possible predictions. The implementation algorithm of the method is as follows:

**Step 1:** Define the initial training set.

**Step 2:** Train the model on the initial training set.

**Step 3:** For each Ag not included in the initial training set:

1. Take the value of model prediction as 0 for each Ab and calculate the loss function value based on this prediction;
2. Take the value of model prediction as 1 for each Ab and calculate the loss function value based on this prediction;
3. Calculate the gradient of the loss function with respect to the last linear layer for the prediction value of 0 and calculate its norm for given Ag and all Abs, then repeat the same for the prediction value of 1;
4. Calculate the distance between the gradient vectors for prediction 0 and prediction 1;
5. Calculate the average distance value for all Abs for given Ag.

**Step 4:** Add the Ag with the largest average distance between gradient vectors for all Abs to the training set.

The **Gradient 0-1 Max** (Grad 0-1 Max) method differs from the Gradient 0-1 Average method only by considering not the average distance value for all Abs, but its maximum value.

The **Gradient to Input Average** (Grad-In Avg) method calculates the gradient of the loss function with respect to the input data: similar to the Gradient 0-1 Average method, the values 0 and 1 are substituted into the loss function, and then the distance between the gradients of the loss function with respect to the input value of the Ab-Ag pair embedding is calculated. Thus, the gradient for each Ab-Ag pair will be a tensor of dimension {*Length of the input amino acid sequence for the Ab-Ag pair*} *x* {*Embedding dimension*}. Next, the vector obtained by summing over the embedding dimension is considered, and for each pair of vectors corresponding to the prediction values 0 and 1, the distance between them is calculated. The Ag with the highest average distance over all Abs is then added to the training set. The **Gradient to Input Max** (Grad-In Max) method considers the maximum distance instead of the average.

In the **Gradient to Model Average** (Grad-Model Avg) method, instead of the loss function, the model prediction is considered as a function of weights: for each Ab-Ag pair, the model prediction for their binding is calculated, and then its gradient is calculated for the last (linear) layer of the model. The Ag with the highest average distance over all Abs is selected. In the **Gradient to Model Max** (Grad-Model Max) method, the maximum gradient norm for the set of Abs is maximized, rather than the average one.

The final method **Gradient Conf. Labels Average** (Grad-Conf Avg) is based on the confounding label approach (Lee and AlRegib, 2020). A confounding label is an artificially introduced label that has not been observed by the model before. The confounding label principle is based on the following observation: if the model is familiar with the input sample, it only needs to learn the patterns between the learned features and the label that is new to it. If the model is not familiar with the sample (and therefore this sample is more informative for the model), it will need to learn new features and then associate them with the labels, which should take more resources. To implement the confounding label method, the model is rebuilt for the multi-classification problem to output two probability values instead of one (one value corresponds to 0, and the other to 1, and together they sum up to 1). The full algorithm for the confounding label method is as follows:

**Step 1:** Define the initial training set.

**Step 2:** Train the multi classification model (for two classes) on the initial training set.

**Step 3:** For each Ag not included in the initial training set:

1. Calculate the value of the loss function (the loss function for the multi classification problem) based on the new confounding label substituted into it for each Ab-Ag pair;
2. Calculate the gradient of the loss function with respect to the last linear layer of the model and calculate its norm for the given Ag and all Abs;
3. Calculate the average value of the gradient norm for all Abs for the given Ag.

**Step 4:** Add the Ag with the highest average gradient norm for all Abs to the training set.

The **Gradient Conf. Labels Max** (Grad-Conf Max) method differs from the Gradient Conf. Labels Average method only in that it maximizes not the average value of the gradient norm for all Abs, but its maximum value.

Thus, many different approaches to measuring model uncertainty for a sample using the model gradient were considered. It is also worth noting that gradient approaches with respect to other model layers except the last one were not considered, which may be of additional interest.

#### 2.2 Diversity based

##### 2.2.1 Clustering methods

The distance-based approach of (Sener and Savarese, 2018) was shown to be effective for an image classification task, but we did not manage to find similar studies in the field of immunology. Our goal was to consistently identify the most informative elements of the sample, which are amino acid sequences, so we implemented an algorithm for selecting the next Ag based on the distance between it and the original training set of Ags. We considered the average, minimum, and maximum distances as the distance between the Ag and the training set of Ags. The Hamming distance, Damerau-Levenshtein distance, and alignment-based distance (Cock et al., 2009) were considered as the distance between pairs of Ags. Hamming distance showed the best results among other distances used. The performance of methods based on minimizing the average Hamming distance and the average alignment-based distance are compared in the article. The result of the method based on minimizing the minimum Hamming distance is also presented.

A brief description of the algorithm steps is presented in Figure 4:

**Figure 4:**
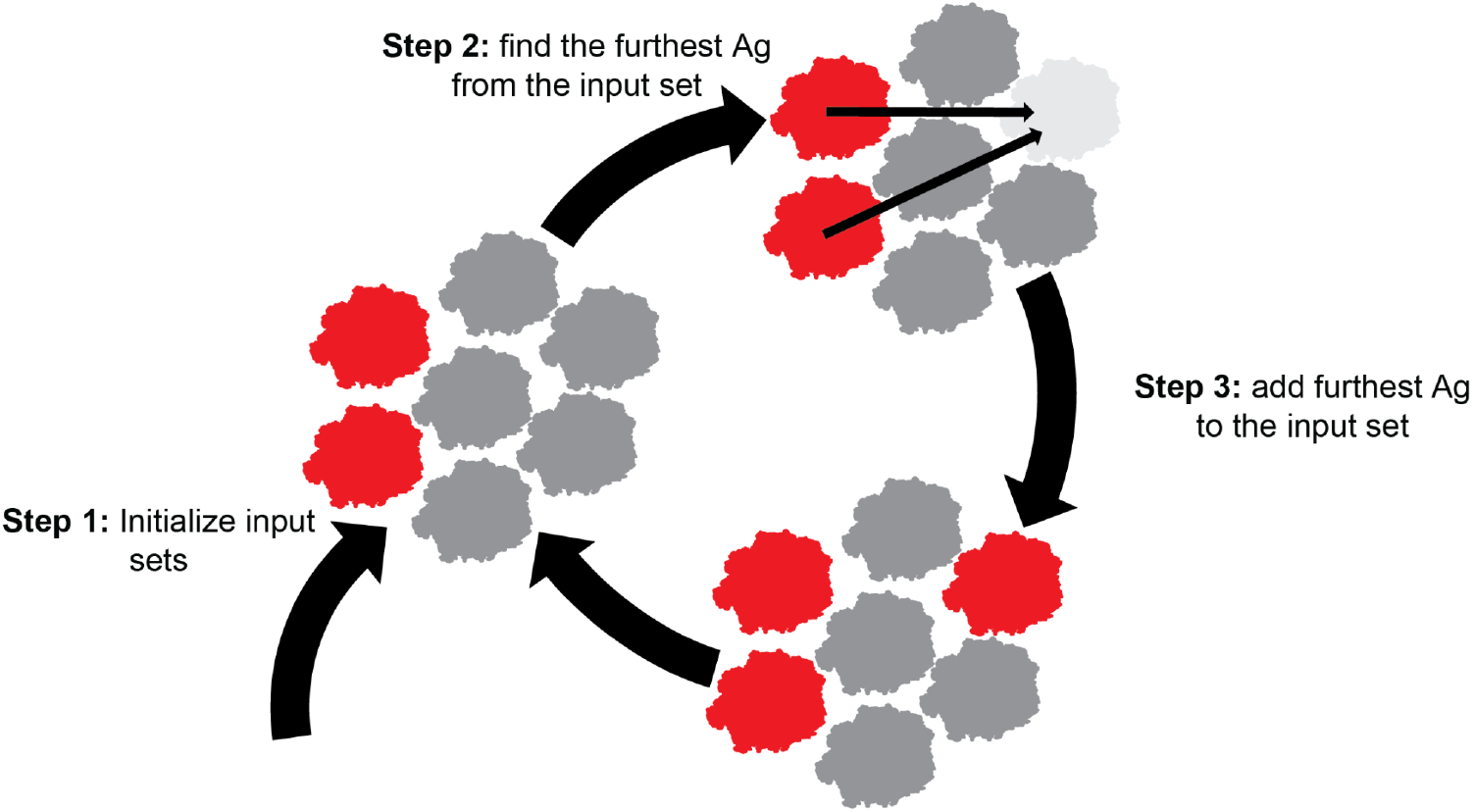
Clustering-based methods select novel Ags based on their similarity to already labeled Ags. Clustering-based methods consist of three steps. Step 1: Initialize input set. Step 2: Find the unlabeled Ag that is furthest from the input set. Step 3: Label the selected Ag and add it to the set of labeled training data.

The algorithm for selecting Ags based on distance between them includes the following steps (Figure 4):

**Step 1:** Define the initial training set.

**Step 2:** For each sample element not included in the initial training set, calculate the distance between it and the training set.

**Step 3:** Add the Ag with the largest distance to the training set.

The algorithm sequentially selects Ags with the largest distance to the initial training set. In the Hamming Average Distance approach, the Ag with the highest mean Hamming distance to all labeled Ags is selected to ensure overall diversity. The Hamming Minimum Distance strategy instead chooses the Ag with the largest smallest Hamming distance to any labeled Ag. For the Alignment Average Distance method, the Ag with the largest mean alignment-based distance to all labeled Ags is selected.

### 3. Packages used

Our computational work relied on the following libraries: PyTorch 2.2.2 (Paszke et al., 2019), NumPy 2.2.1 (Harris et al., 2020), pandas 2.2.3 (McKinney, 2010) and SciPy 1.6.1 (SciPy 1.0 Contributors et al., 2020). Graphics were generated with Matplotlib 3.10.0 (Hunter, 2007) and Seaborn 0.13.2 (Waskom, 2021).

## Availability of data and materials

All code is available at https://github.com/csi-greifflab/AbAgAL.

## Author contributions

DB and RF designed the methodology. RF generated data. DB, RF, SK and DW developed and evaluated the active learning strategies. DB, RF and SK wrote the manuscript. PR advised on data generation. All authors edited and approved the final manuscript.

## Funding

This work was supported by ARCAID (https://www.arcaid-h2020.eu), which received funding from the European Union’s Horizon 2020 research and innovation programme under the Marie Skłodowska-Curie grant agreement No 847551. The Leona M. and Harry B. Helmsley Charitable Trust (#2019PG-T1D011, to VG), UiO World-Leading Research Community (to VG), UiO: LifeScience Convergence Environment Immunolingo (to VG and GKS), EU Horizon 2020 iReceptorplus (#825821) (to VG), a Norwegian Cancer Society Grant (#215817, to VG), Research Council of Norway projects (#300740, #331890 to VG), a Research Council of Norway IKTPLUSS project (#311341, to VG and GKS), and Stiftelsen Kristian Gerhard Jebsen (K.G. Jebsen Coeliac Disease Research Centre) (to GKS). This project has received funding (to VG) from the Innovative Medicines Initiative 2 Joint Undertaking under grant agreement No 101007799 (Inno4Vac). This Joint Undertaking receives support from the European Union’s Horizon 2020 research and innovation programme and EFPIA. This communication reflects the author’s view and neither IMI nor the European Union, EFPIA, or any Associated Partners are responsible for any use that may be made of the information contained therein. Funded by the European Union (ERC, AB-AG-INTERACT, 101125630, to VG).

## Competing interests

V.G. declares advisory board positions in aiNET GmbH, Enpicom B.V, Absci, Omniscope, and Diagonal Therapeutics. V.G. is a consultant for Adaptive Biosystems, Specifica Inc, Roche/Genentech, immunai, LabGenius, and FairJourney Biologics.

PR declares employment by F. Hoffmann-La Roche AG.

## Acknowledgments

We thank Prof. dr. Antoine van Kampen and Dr. ir. Perry Moerland for their contributions to the project and valuable discussions.

## Notes

https://github.com/csi-greifflab/AbAgAL

### Summary of Updates

Improvements to Figure 1; corrected caption for Figure 3

## References

Ash, J.T., Zhang, C., Krishnamurthy, A., Langford, J., Agarwal, A., 2019. Deep Batch Active Learning by Diverse, Uncertain Gradient Lower Bounds. Presented at the International Conference on Learning Representations.

Buntz, B., 2024. Genentech’s lab in the loop aims to tap the power of quantity for quality drug discovery. URL https://www.drugdiscoverytrends.com/genentech-ai-lab-in-the-loop-drug-discovery/

Burley, S.K., Bhikadiya, C., Bi, C., Bittrich, S., Chao, H., Chen, L., Craig, P.A., Crichlow, G.V., Dalenberg, K., Duarte, J.M., Dutta, S., Fayazi, M., Feng, Z., Flatt, J.W., Ganesan, S., Ghosh, S., Goodsell, D.S., Green, R.K., Guranovic, V., Henry, J., Hudson, B.P., Khokhriakov, I., Lawson, C.L., Liang, Y., Lowe, R., Peisach, E., Persikova, I., Piehl, D.W., Rose, Y., Sali, A., Segura, J., Sekharan, M., Shao, C., Vallat, B., Voigt, M., Webb, B., Westbrook, J.D., Whetstone, S., Young, J.Y., Zalevsky, A., Zardecki, C., 2023. RCSB Protein Data Bank (RCSB.org): delivery of experimentally-determined PDB structures alongside one million computed structure models of proteins from artificial intelligence/machine learning. Nucleic Acids Res 51, D488–D508. 10.1093/nar/gkac1077

Chinery, L., Hummer, A.M., Mehta, B.B., Akbar, R., Rawat, P., Slabodkin, A., Quy, K.L., Lund-Johansen, F., Greiff, V., Jeliazkov, J.R., Deane, C.M., 2024. Baselining the Buzz Trastuzumab-HER2 Affinity, and Beyond. 10.1101/2024.03.26.586756

Cock, P.J.A., Antao, T., Chang, J.T., Chapman, B.A., Cox, C.J., Dalke, A., Friedberg, I., Hamelryck, T., Kauff, F., Wilczynski, B., De Hoon, M.J.L., 2009. Biopython: freely available Python tools for computational molecular biology and bioinformatics. Bioinformatics 25, 1422–1423. 10.1093/bioinformatics/btp163

Cohn, D., Atlas, L., Ladner, R., 1994. Improving generalization with active learning. Mach Learn 15, 201–221. 10.1007/BF00993277

Dauparas, J., Anishchenko, I., Bennett, N., Bai, H., Ragotte, R.J., Milles, L.F., Wicky, B.I.M., Courbet, A., De Haas, R.J., Bethel, N., Leung, P.J.Y., Huddy, T.F., Pellock, S., Tischer, D., Chan, F., Koepnick, B., Nguyen, H., Kang, A., Sankaran, B., Bera, A.K., King, N.P., Baker, D., 2022. Robust deep learning–based protein sequence design using ProteinMPNN. Science 378, 49–56. 10.1126/science.add2187

Ehling, R.A., Minot, M., Overath, M.D., Sheward, D.J., Han, J., Gao, B., Taft, J.M., Pertseva, M., Weber, C.R., Frei, L., Bikias, T., Murrell, B., Reddy, S.T., 2024. Synthetic coevolution reveals adaptive mutational trajectories of neutralizing antibodies and SARS-CoV-2. 10.1101/2024.03.28.587189

Engelhart, E., Emerson, R., Shing, L., Lennartz, C., Guion, D., Kelley, M., Lin, C., Lopez, R., Younger, D., Walsh, M.E., 2022a. A dataset comprised of binding interactions for 104,972 antibodies against a SARS-CoV-2 peptide. Sci Data 9, 653. 10.1038/s41597-022-01779-4

Engelhart, E., Lopez, R., Emerson, R., Lin, C., Shikany, C., Guion, D., Kelley, M., Younger, D., 2022b. Massively multiplexed affinity characterization of therapeutic antibodies against SARS-CoV-2 variants. Antib Ther 5, 130–137. 10.1093/abt/tbac011

Gessner, A., Ober, S.W., Vickery, O., Oglić, D., Uçar, T., 2024. Active learning for affinity prediction of antibodies. 10.48550/ARXIV.2406.07263

Graves, J., Byerly, J., Priego, E., Makkapati, N., Parish, S., Medellin, B., Berrondo, M., 2020. A Review of Deep Learning Methods for Antibodies. Antibodies 9, 12. 10.3390/antib9020012

Harris, C.R., Millman, K.J., Van Der Walt, S.J., Gommers, R., Virtanen, P., Cournapeau, D., Wieser, E., Taylor, J., Berg, S., Smith, N.J., Kern, R., Picus, M., Hoyer, S., Van Kerkwijk, M.H., Brett, M., Haldane, A., Del Río, J.F., Wiebe, M., Peterson, P., Gérard-Marchant, P., Sheppard, K., Reddy, T., Weckesser, W., Abbasi, H., Gohlke, C., Oliphant, T.E., 2020. Array programming with NumPy. Nature 585, 357–362. 10.1038/s41586-020-2649-2

Henikoff, S., Henikoff, J.G., 1992. Amino acid substitution matrices from protein blocks. Proc. Natl. Acad. Sci. U.S.A. 89, 10915–10919. 10.1073/pnas.89.22.10915

Huang, Y., Zhang, Z., Zhou, Y., 2022. AbAgIntPre: A deep learning method for predicting antibody-antigen interactions based on sequence information. Front. Immunol. 13, 1053617. 10.3389/fimmu.2022.1053617

Hunter, J.D., 2007. Matplotlib: A 2D Graphics Environment. Comput. Sci. Eng. 9, 90–95. 10.1109/MCSE.2007.55

Jin, R., Ye, Q., Wang, J., Cao, Z., Jiang, D., Wang, T., Kang, Y., Xu, W., Hsieh, C.-Y., Hou, T., 2024. AttABseq: an attention-based deep learning prediction method for antigen–antibody binding affinity changes based on protein sequences. Briefings in Bioinformatics 25. 10.1093/bib/bbae304

Khan, A., Cowen-Rivers, A.I., Grosnit, A., Deik, D.-G.-X., Robert, P.A., Greiff, V., Smorodina, E., Rawat, P., Akbar, R., Dreczkowski, K., Tutunov, R., Bou-Ammar, D., Wang, J., Storkey, A., Bou-Ammar, H., 2023. Toward real-world automated antibody design with combinatorial Bayesian optimization. Cell Reports Methods 3, 100374. 10.1016/j.crmeth.2022.100374

Lee, J., AlRegib, G., 2020. Gradients as a Measure of Uncertainty in Neural Networks, in: 2020 IEEE International Conference on Image Processing (ICIP). Presented at the 2020 IEEE International Conference on Image Processing (ICIP), pp. 2416–2420. 10.1109/ICIP40778.2020.9190679

Li, L., Gupta, E., Spaeth, J., Shing, L., Jaimes, R., Engelhart, E., Lopez, R., Caceres, R.S., Bepler, T., Walsh, M.E., 2023. Machine learning optimization of candidate antibody yields highly diverse sub-nanomolar affinity antibody libraries. Nat Commun 14, 3454. 10.1038/s41467-023-39022-2

Lin, Z., Akin, H., Rao, R., Hie, B., Zhu, Z., Lu, W., Smetanin, N., Verkuil, R., Kabeli, O., Shmueli, Y., Dos Santos Costa, A., Fazel-Zarandi, M., Sercu, T., Candido, S., Rives, A., 2023. Evolutionary-scale prediction of atomic-level protein structure with a language model. Science 379, 1123–1130. 10.1126/science.ade2574

Lundberg, S.M., Lee, S.-I., 2017. A Unified Approach to Interpreting Model Predictions, in: Advances in Neural Information Processing Systems. Curran Associates, Inc.

Lyu, X., Zhao, Q., Hui, J., Wang, T., Lin, M., Wang, K., Zhang, J., Shentu, J., Dalby, P.A., Zhang, H., Liu, B., 2022. The global landscape of approved antibody therapies. Antibody Therapeutics 5, 233–257. 10.1093/abt/tbac021

Maraval, A., Zimmer, M., Grosnit, A., Bou Ammar, H., 2023. End-to-End Meta-Bayesian Optimisation with Transformer Neural Processes. Advances in Neural Information Processing Systems 36, 11246–11260.

Mason, D.M., Reddy, S.T., 2024. Predicting adaptive immune receptor specificities by machine learning is a data generation problem. Cell Systems 15, 1190–1197. 10.1016/j.cels.2024.11.008

McKinney, W., 2010. Data Structures for Statistical Computing in Python. Presented at the Python in Science Conference, Austin, Texas, pp. 56–61. 10.25080/Majora-92bf1922-00a

Murphy, K.M., Weaver, C., 2017. Janeway’s immunobiology, 9th edition. ed. GS, Garland Science, Taylor & Francis Group, New York London.

Nelson, P.N., 2000. Demystified …: Monoclonal antibodies. Molecular Pathology 53, 111–117. 10.1136/mp.53.3.111

O’Donnell, T.J., Kanduri, C., Isacchini, G., Limenitakis, J.P., Brachman, R.A., Alvarez, R.A., Haff, I.H., Sandve, G.K., Greiff, V., 2024. Reading the repertoire: Progress in adaptive immune receptor analysis using machine learning. Cell Systems 15, 1168–1189. 10.1016/j.cels.2024.11.006

Olsen, T.H., Moal, I.H., Deane, C.M., 2022. AbLang: an antibody language model for completing antibody sequences. Bioinformatics Advances 2, vbac046. 10.1093/bioadv/vbac046

Paszke, A., Gross, S., Massa, F., Lerer, A., Bradbury, J., Chanan, G., Killeen, T., Lin, Z., Gimelshein, N., Antiga, L., Desmaison, A., Köpf, A., Yang, E., DeVito, Z., Raison, M., Tejani, A., Chilamkurthy, S., Steiner, B., Fang, L., Bai, J., Chintala, S., 2019. PyTorch: An Imperative Style, High-Performance Deep Learning Library. 10.48550/ARXIV.1912.01703

Pochtovyi, A.A., Kustova, D.D., Siniavin, A.E., Dolzhikova, I.V., Shidlovskaya, E.V., Shpakova, O.G., Vasilchenko, L.A., Glavatskaya, A.A., Kuznetsova, N.A., Iliukhina, A.A., Shelkov, A.Y., Grinkevich, O.M., Komarov, A.G., Logunov, D.Y., Gushchin, V.A., Gintsburg, A.L., 2023. In Vitro Efficacy of Antivirals and Monoclonal Antibodies against SARS-CoV-2 Omicron Lineages XBB.1.9.1, XBB.1.9.3, XBB.1.5, XBB.1.16, XBB.2.4, BQ.1.1.45, CH.1.1, and CL.1. Vaccines (Basel) 11, 1533. 10.3390/vaccines11101533

Ragonnet-Cronin, M., Nutalai, R., Huo, J., Dijokaite-Guraliuc, A., Das, R., Tuekprakhon, A., Supasa, P., Liu, C., Selvaraj, M., Groves, N., Hartman, H., Ellaby, N., Mark Sutton, J., Bahar, M.W., Zhou, D., Fry, E., Ren, J., Brown, C., Klenerman, P., Dunachie, S.J., Mongkolsapaya, J., Hopkins, S., Chand, M., Stuart, D.I., Screaton, G.R., Rokadiya, S., 2023. Generation of SARS-CoV-2 escape mutations by monoclonal antibody therapy. Nat Commun 14, 3334. 10.1038/s41467-023-37826-w

Rainforth, T., Foster, A., Ivanova, D.R., Smith, F.B., 2023. Modern Bayesian Experimental Design. 10.48550/arXiv.2302.14545

Ren, P., Xiao, Y., Chang, X., Huang, P.-Y., Li, Z., Gupta, B.B., Chen, X., Wang, X., 2021. A Survey of Deep Active Learning. ACM Comput. Surv. 54, 180:1-180:40. 10.1145/3472291

Ribeiro, M.T., Singh, S., Guestrin, C., 2016. “Why Should I Trust You?”: Explaining the Predictions of Any Classifier, in: Proceedings of the 22nd ACM SIGKDD International Conference on Knowledge Discovery and Data Mining, KDD ’16. Association for Computing Machinery, New York, NY, USA, pp. 1135–1144. 10.1145/2939672.2939778

Robert, P.A., Akbar, R., Frank, R., Pavlović, M., Widrich, M., Snapkov, I., Slabodkin, A., Chernigovskaya, M., Scheffer, L., Smorodina, E., Rawat, P., Mehta, B.B., Vu, M.H., Mathisen, I.F., Prósz, A., Abram, K., Olar, A., Miho, E., Haug, D.T.T., Lund-Johansen, F., Hochreiter, S., Haff, I.H., Klambauer, G., Sandve, G.K., Greiff, V., 2022. Unconstrained generation of synthetic antibody-antigen structures to guide machine learning methodology for antibody specificity prediction. Nat Comput Sci 2, 845–865. 10.1038/s43588-022-00372-4

SciPy 1.0 Contributors, Virtanen, P., Gommers, R., Oliphant, T.E., Haberland, M., Reddy, T., Cournapeau, D., Burovski, E., Peterson, P., Weckesser, W., Bright, J., Van Der Walt, S.J., Brett, M., Wilson, J., Millman, K.J., Mayorov, N., Nelson, A.R.J., Jones, E., Kern, R., Larson, E., Carey, C.J., Polat, İ., Feng, Y., Moore, E.W., VanderPlas, J., Laxalde, D., Perktold, J., Cimrman, R., Henriksen, I., Quintero, E.A., Harris, C.R., Archibald, A.M., Ribeiro, A.H., Pedregosa, F., Van Mulbregt, P., 2020. Author Correction: SciPy 1.0: fundamental algorithms for scientific computing in Python. Nat Methods 17, 352–352. 10.1038/s41592-020-0772-5

Sener, O., Savarese, S., 2018. Active Learning for Convolutional Neural Networks: A Core-Set Approach. Presented at the International Conference on Learning Representations.

Seo, S., Kwak, M.W., Kang, E., Kim, C., Park, E., Kang, T.H., Kim, J., 2022. Accelerating Antibody Design with Active Learning. 10.1101/2022.09.12.507690

Settles, B., 2009. Active Learning Literature Survey (Technical Report). University of Wisconsin-Madison Department of Computer Sciences.

Seung, H.S., Opper, M., Sompolinsky, H., 1992. Query by committee, in: Proceedings of the Fifth Annual Workshop on Computational Learning Theory, COLT ’92. Association for Computing Machinery, New York, NY, USA, pp. 287–294. 10.1145/130385.130417

Taft, J.M., Weber, C.R., Gao, B., Ehling, R.A., Han, J., Frei, L., Metcalfe, S.W., Overath, M.D., Yermanos, A., Kelton, W., Reddy, S.T., 2022. Deep mutational learning predicts ACE2 binding and antibody escape to combinatorial mutations in the SARS-CoV-2 receptor-binding domain. Cell 185, 4008-4022.e14. 10.1016/j.cell.2022.08.024

Tonegawa, S., 1983. Somatic generation of antibody diversity. Nature 302, 575–581. 10.1038/302575a0

Towers, S., Kalisz, A., Robert, P.A., Higueruelo, A., Vianello, F., Tsai, M.-H.C., Steel, H., Foerster, J.N., 2024. Opponent Shaping for Antibody Development. 10.48550/arXiv.2409.10588

Waskom, M., 2021. seaborn: statistical data visualization. JOSS 6, 3021. 10.21105/joss.03021

