## Supplementary table 1 for "Active learning for improving out-of-distribution lab-in-the-loop experimental design"

### Hamming Average Distance, QBC and Gradient-Based Methods show statistically significant performance

The relative improvement in AUC, referred to as relative difference, is calculated by comparing the AUC performance of a specific method against the Random baseline. This is expressed as a percentage increase or decrease.

$$relative\ difference = mean(method\ AUC - Random\ AUC)\ /\ mean(Random\ AUC)$$

Across the different test scenarios (Test, TestSharedAG, and TestSharedAB), as shown in Table 1, the performance of methods varies. Notably, QBC consistently shows statistically significant improvements over the random baseline in multiple scenarios. The Hamming Average Distance and Gradient on Last Layer (both Max and Average) methods exhibit mixed results, with some configurations failing to show significant improvements. Such findings suggest that while certain methods can enhance performance, their efficacy may be context-dependent, necessitating further investigation to optimize their application.

| test | method | p-value | significance | CI lower | CI upper | relative difference |
|---|---|---|---|---|---|---|
| Test | Alignments average distance | 1,000 | FALSE | -0,475 | inf | 0,044% |
| | Gradient to input (average) | 1,000 | FALSE | -0,868 | inf | -0,516% |
| | Gradient to input (max) | 1,000 | FALSE | -0,835 | inf | -0,450% |
| | Gradient to model (average) | 1,000 | FALSE | -1,416 | inf | -1,263% |
| | Gradient to model (max) | 1,000 | FALSE | -1,387 | inf | -1,222% |
| | Gradient 0-1 (average) | 1,000 | FALSE | -1,459 | inf | -1,322% |
| | Gradient 0-1 (max) | 1,000 | FALSE | -1,355 | inf | -1,184% |
| | Gradient on last layer (average) | 1,000 | FALSE | -0,334 | inf | 0,217% |
| | Gradient conf. labels (average) | 1,000 | FALSE | -1,161 | inf | -0,940% |
| | Gradient conf. labels (max) | 1,000 | FALSE | -1,199 | inf | -0,958% |
| | Gradient on last layer (max) | 0,075 | FALSE | -0,101 | inf | 0,534% |
| | Hamming average distance | 0,000 | TRUE | 0,873 | inf | 1,795% |
| | Hamming min distance | 1,000 | FALSE | -2,120 | inf | -2,119% |
| | Query-by-Committee | 0,000 | TRUE | 0,136 | inf | 0,766% |
| TestSharedAG | Alignments average distance | 1,000 | FALSE | -0,245 | inf | 0,080% |
| | Gradient to input (average) | 1,000 | FALSE | -0,913 | inf | -0,758% |
| | Gradient to input (max) | 1,000 | FALSE | -0,778 | inf | -0,591% |

| | | | | | | |
|---|---|---|---|---|---|---|
| | Gradient to model (average) | 1,000 | FALSE | -1,625 | inf | -1,681% |
| | Gradient to model (max) | 1,000 | FALSE | -1,570 | inf | -1,593% |
| | Gradient 0-1 (average) | 1,000 | FALSE | -1,540 | inf | -1,583% |
| | Gradient 0-1 (max) | 1,000 | FALSE | -1,576 | inf | -1,602% |
| | Gradient on last layer (average) | 0,002 | TRUE | 0,046 | inf | 0,449% |
| | Gradient conf. labels (average) | 1,000 | FALSE | -1,343 | inf | -1,324% |
| | Gradient conf. labels (max) | 1,000 | FALSE | -1,298 | inf | -1,257% |
| | Gradient on last layer (max) | 0,000 | TRUE | 0,144 | inf | 0,574% |
| | Hamming average distance | 0,024 | FALSE | -0,026 | inf | 0,357% |
| | Hamming min distance | 1,000 | FALSE | -1,468 | inf | -1,417% |
| | Query-by-Committee | 0,006 | TRUE | 0,014 | inf | 0,405% |
| TestSharedAB | Alignments average distance | 1,000 | FALSE | -0,353 | inf | 0,102% |
| | Gradient to input (average) | 1,000 | FALSE | -0,702 | inf | -0,322% |
| | Gradient to input (max) | 1,000 | FALSE | -0,795 | inf | -0,411% |
| | Gradient to model (average) | 1,000 | FALSE | -1,402 | inf | -1,203% |
| | Gradient to model (max) | 1,000 | FALSE | -1,395 | inf | -1,194% |
| | Gradient 0-1 (average) | 1,000 | FALSE | -1,470 | inf | -1,267% |
| | Gradient 0-1 (max) | 1,000 | FALSE | -1,387 | inf | -1,183% |
| | Gradient on last layer (average) | 1,000 | FALSE | -0,296 | inf | 0,177% |
| | Gradient conf. labels (average) | 1,000 | FALSE | -1,399 | inf | -1,162% |
| | Gradient conf. labels (max) | 1,000 | FALSE | -1,417 | inf | -1,170% |
| | Gradient on last layer (max) | 0,043 | FALSE | -0,065 | inf | 0,471% |
| | Hamming average distance | 0,000 | TRUE | 0,663 | inf | 1,309% |
| | Hamming min distance | 1,000 | FALSE | -1,790 | inf | -1,582% |
| | Query-by-Committee | 0,000 | TRUE | 0,266 | inf | 0,777% |

**Supplementary table 1: Statistical comparison of AUCs between baseline (random order of added Ags) and various methods.** Table presents the results of the paired Student's t-test comparing the ROC AUC of the Random method to those of various alternative methods, with a Bonferroni correction applied. The null hypothesis for the t-test states that the Random AUC is greater than or equal to the AUC of each alternative method. Statistically significant results are marked in green.
