## Supplementary figure 1 for "Active learning for improving out-of-distribution lab-in-the-loop experimental design"

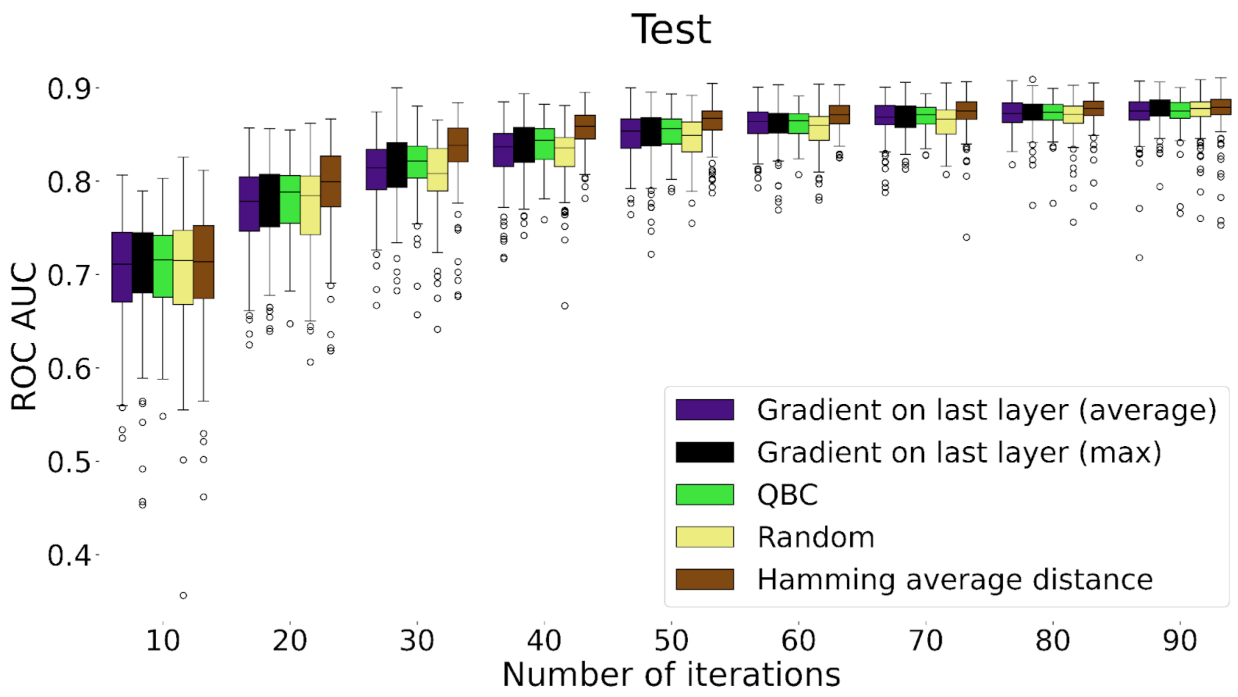


**Supplementary figure 1: ROC AUC performance Box Plots for Test scenario.** The figure presents the ROC AUC values of the four outperforming AL approaches along with the random baseline. Box plots correspond to every 10th iteration, with each data point representing one of 200 experimental repetitions.
